# Adaptations of energy metabolism in cetaceans have consequences for their response to foraging disruption

**DOI:** 10.1101/709154

**Authors:** Davina Derous, Jagajjit Sahu, Alex Douglas, David Lusseau, Marius Wenzel

## Abstract

Cetaceans have varied their anatomical structure, physiology and metabolism to adapt to the challenges of aquatic life. Key to this change is the deposition of blubber. This adipose tissue plays a significant regulatory and signaling role in mammalian metabolism. As foraging disruption by human activities is emerging as a key conservation threat for cetaceans, we need to understand how selection for aquatic life might have altered key nutrient sensing pathways associated with adipose signaling. We compared selection pressure on those energy metabolism biological pathways by contrasting the rate of substitution observed in genes associated with them in cetacean and artiodactyl genomes. We then estimated the likely consequence of these selection pressures for pathway functions. Here we show that genes involved in the insulin, mTOR, SIRT and NF-κB pathways were under significant positive selection in cetaceans compared to their terrestrial sister taxon. Our results suggest these genes may have been positively selected to adapt to a glucose-poor diet and it is unlikely that fat depots signaling function in the same manner as in terrestrial mammals. Secondary adaptation to life in water significantly affected functions in nutrient sensing pathways in cetaceans. Insulin is not likely to play the same role in energy balance as it does in terrestrial mammals and adiposity is not likely to have the deleterious health consequences it has in terrestrial mammals. The physiological ecology of cetacean fat deposition, and therefore its value as a condition index, needs to be interpreted in this evolutionary context.

## Introduction

Cetaceans are mammals that transitioned from a terrestrial to an aquatic lifestyle approximately 53–56 million years ago by adapting their anatomical structure, physiology and metabolism. These critical morphological and physiological adaptations ensured the maintenance of body temperature and energy reserves (Parry, 1949; Scholander, Walters, Hock, & Irving, 1950). For example, this included a thickening of the blubber (i.e. adipose tissue equivalent) to provide thermal insulation, to deal with more sporadic foraging opportunities, and to support locomotion (Vasseur & Yodzis, 2004; T M Williams, Friedl, & Haun, 1993; Terrie M. Williams, Haun, Davis, Fuiman, & Kohin, 2001). Crucially, these adaptations impact the ways in which individuals decide to invest in reproduction and define their abilities to survive under varied environmental pressures, in particular nutrient availability. Human-caused perturbations, such as shipping, tourism, naval activities, coastal urbanization and offshore energy development, can perturb environmental nutrient levels and affect cetacean foraging abilities. These factors are becoming a pervasive and prevalent threat to many cetacean species (Pirotta et al., 2018) and are a key priority in cetacean conservation policy (eg. National Academies of Sciences, Engineering, 2017).

Cetaceans detect fluctuations in environmental nutrient levels by nutrient sensing pathways and some of these are evolutionary conserved across species (Chantranupong, Wolfson, & Sabatini, 2015). These pathways detect intracellular and extracellular levels of sugar, amino acids and lipids and their surrogate metabolites. Nutrients can trigger the release of several hormones, which induce coherent responses in several pathways involved in regulating metabolism. Unsurprisingly, a large portion of positively selected genes in cetaceans are involved in energy metabolism (Nery, González, & Opazo, 2013). Bottlenose dolphins (*Tursiops truncatus*) show insulin resistance likely caused by early metabolic shifts in substrate utilization as the species shifted from a terrestrial high carbohydrate diet to a marine high protein diet (Wang et al., 2016). Dolphin diet has a high fat and protein content and is almost devoid of carbohydrates (Wells et al., 2013). Fasted healthy bottlenose dolphins (*Tursiops truncatus*) have elevated fasting plasma glucose concentration that are similar to diabetic humans (S. Venn-Watson, Carlin, & Ridgway, 2011; S. K. Venn-Watson & Ridgway, 2007).. During fasting, metabolism is believed to be primarily fueled by large adipose stores (i.e. blubber) that are mobilized in response to insulin suppression (Duncan, Ahmadian, Jaworski, Sarkadi-Nagy, & Sul, 2007).

This fasting response is expected to take place when foraging is disrupted in the wild by human activities. Regulatory genes related to lipolysis are positively selected in cetacean-specific lineages but not in terrestrial mammals (Wang et al., 2015). Specifically, genes related to triacylglycerol (TAG) metabolism were suggested to play an essential role in the secondary adaption of cetaceans to aquatic life (Wang et al., 2015). The processes of lipid deposition and utilization is regulated by the gene leptin (LEP) (Duncan et al., 2007). Recent work shows that in bowhead whales (*Balaena mysticetus*) and belugas (*Delphinapterus leucas*), the regulation of LEP and lipolysis is adapted to seasonal cycles of blubber deposition and utilization (Ball et al., 2017). Although adipose tissue biology of terrestrial mammals show a similarity to the functioning of cetacean blubber, some differences in key genes have been identified (Ball et al., 2017). These changes have therefore the scope to alter the way individual take biological decisions about demographic contributions, particularly reproduction, given their energetic metabolic state. We need to place gene selection in the context of the biological pathways in which they are involved to contextualize those changes and understand the potential demographic consequences of foraging disruption when the environmental nutrient levels of cetaceans are perturbed. Here, we aimed to determine whether the selective pressure from secondary adaptations to life in water led to a change in the key nutrient signaling pathways. We aim to predict whether these changes are likely to affect biological functions.

## Material and Methods

To understand the evolutionary-driven changes in the regulation of metabolic processes in cetaceans, we took a targeted approach and focused on 6 signaling pathways: the p53 signaling pathway, the insulin signaling pathway, the mTOR signaling pathway, the leptin signaling pathway, the NF-κB signaling pathway and the SIRT signaling pathway. A total of 532 genes involved in these pathways were obtained from the Kyoto Encyclopedia of Genes and Genomes (KEGG) website (http://www.genome.jp/kegg/) and from the Ingenuity Pathway Analysis (IPA) program (version 2000-2019, Ingenuity Systems, www.ingenuity.com).

### Analysis of positive gene selection and amino-acid substitutions

We obtained the full amino-acid sequences from 532 human KEGG proteins included in our pathways of interest and downloaded from *NCBI* the genome assemblies of human, mouse, 16 cetacean species and 37 artiodactyl species (Table S1). We aligned the KEGG proteins to all genomes using *EXONERATE* 2.2.0 (Slater & Birney, 2005) with the *protein2genome* model to allow for spliced alignments across introns. For each genome, the single match with the highest alignment score was retained and the nucleotide sequence of the match was extracted. If multiple best matches with the same score were present, one match was chosen at random. Sequences were codon-aligned with guidance from corresponding amino-acid alignments and allowing for frame-shifts using *MACSE* 2.03 (Ranwez, Douzery, Cambon, Chantret, & Delsuc, 2018), and maximum-likelihood gene trees were inferred in *IQTREE* 1.6.8 (Nguyen, Schmidt, Von Haeseler, & Minh, 2015) with automatic selection of nucleotide substitution models. A species tree was then inferred from all 532 gene trees using *ASTRAL-III* 5.6.3 (Zhang, Rabiee, Sayyari, & Mirarab, 2018) and rooted at the two outgroups human and mouse.

Codon sequence evolution was modelled in the *codeml* program of *PAML* 4.9f (Yang, 2007), using the species tree with a trifurcated root (=derooted). To minimize the impact of missing data, all codons with more than 20 % missing data were removed from the alignments using *TRIMAL* 1.4 (Capella-Gutiérrez, Silla-Martínez, & Gabaldón, 2009). Three types of models were run per alignment. First, the null model estimated a single dN:dS ratio (ω) for the entire alignment. Second, the branch model (model = 2; NSsites = 0) estimated a single ω for all cetacean lineages (foreground) and a second ω for all other lineages (background). Third, the branch-site model (model = 2; NSsites = 2) estimated different ω among codons within the foreground and background branches. The branch models and branch-site models were each run twice, either allowing the foreground ω to vary freely or fixing ω at 1 (=neutral evolution). The statistical significance of the free ω estimates of interest was obtained via likelihood-ratio tests carried out in *R* 3.4.0 (R Core Team, 2014). Both the free branch model and the free branch-site model were contrasted with their corresponding neutral models and with the null model by comparing twice the difference in likelihood (2ΔL) of the models against a *Chi*-square distribution with one degree of freedom. *P*-values were corrected for multiple testing across all genes within each type of contrast using the false-discovery-rate method (Benjamini & Hochberg, 1995).

From all branch-site models with free ω that fitted significantly better than the neutral branch-site model and the null model, we identified the specific codons under positive selection in the cetacean lineage using the Bayes Empirical Bayes (BEB) method (Yang, 2007). We then examined whether the amino-acid substitutions in the cetacean lineage at these sites would have detrimental effects on the function of the protein using *PROVEAN* 1.1.5 (Choi, Sims, Murphy, Miller, & Chan, 2012). For each protein, the *Homo sapiens* sequence was used as the reference sequence, and the single most frequent alternative amino acid observed among the 16 cetacean genomes was queried.

### Pathway level comparison

The value of statistically significant likelihood-ratio test statistics (2ΔL) between the positive and neutral branch-models were used as a measure to visualize the interactions of the positive gene selection at a pathway level using IPA signaling pathways for each of the target pathway (version 2000-2019, Ingenuity Systems, www.ingenuity.com). IPA did not have a prebuilt sirtuin signaling pathway. We therefore manually constructed this pathway based on the summarized data by Nakagawa and Guarente (2011) (Nakagawa & Guarente, 2011).

### Prediction of functional effects

Deleterious amino acid substitutions (PROVEAN scores <= -2.5) in positively selected codons were visualized using IPA and possible downstream effects in the pathways based on the damaged protein were predicted using the Molecule Activity Predictor (MAP) function in IPA.

## Results

### Strength of selective pressure on genes involved in nutrient sensing pathways

The species tree inferred from all gene trees (Fig 1) was consistent with published cetacean and artiodactylan phylogenies (Zurano et al., 2019). Alignment-wide ω estimates from the PAML null models were <1 for all but one gene (median: 0.08; mean: 0.12), consistent with a baseline of strong purifying selection on protein function across all taxa and all codons (Fig 2A). Contrasting the cetacean group with all other taxa using branch models revealed that 224 out of 532 genes (42.1 %) departed significantly (q ≤ 0.05) from neutral codon evolution (ω ≠ 1), but all of these genes were under purifying selection (ω <1) instead of positive selection (ω >1). Only seven genes were candidates for positive selection (ω >1), but none of these estimates were significant (Fig 2B). In contrast, branch-site models revealed significant (q ≤ 0.05) positive selection (ω >1; median: 9.67; mean: 39.64) in the cetacean group on a small subset of codons in 133 of 532 genes (25 %) (Fig 2C).

**Figure 1.**
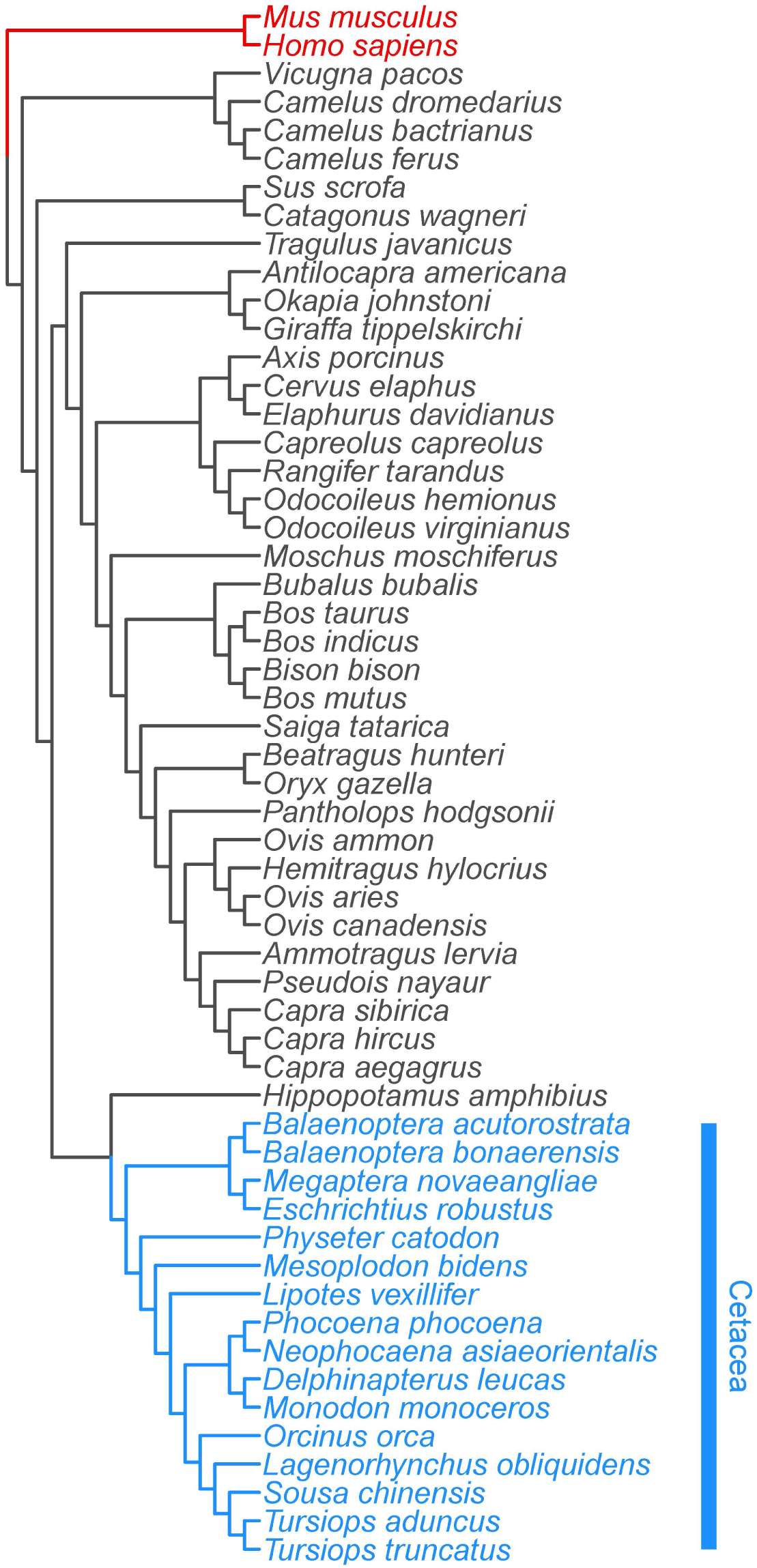
Species cladogram of the cetacea ingroup (blue) and artiodactyla, human and mouse outgroups (black and red), derived from 532 gene alignments.

**Figure 2:**
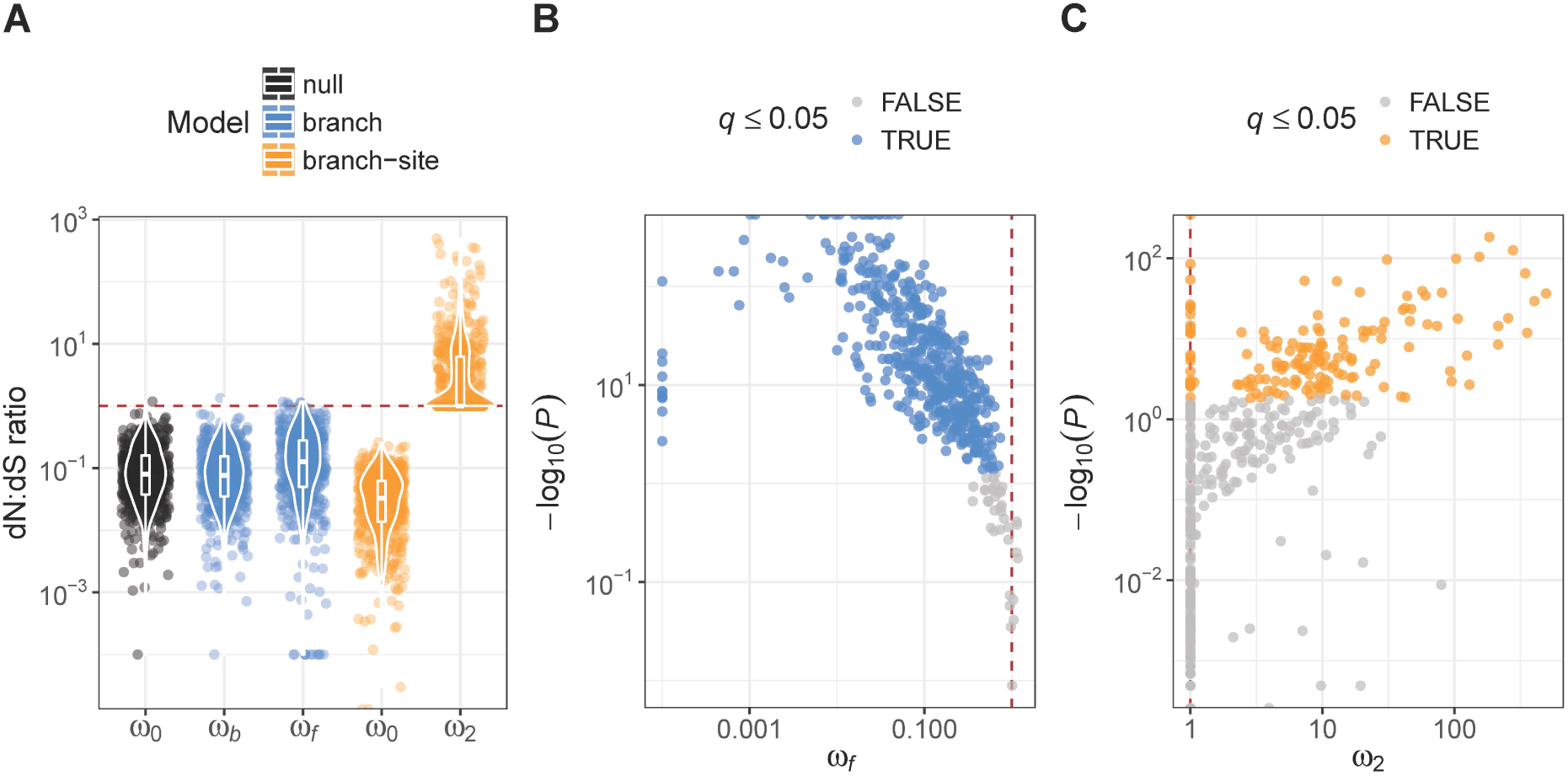
Ratios of non-synonymous vs. synonymous nucleotide substitution rates (dN:dS ratios; ω) in 532 genes estimated from codon evolution models in PAML. A) baseline estimates for whole alignments from null models (ω0), estimates for foreground (cetacea; ωf) and background (all others; ωb) branches from branch models, and estimates for codons under purifying (ω0) and positive (ω2) selection in foreground branch (cetacea) from branch-site models. B) Foreground dN:dS ratio (ωf) and statistical significance (*P*-value) from likelihood-ratio tests between free-ratio branch models and neutral branch models. Significant tests after FDR correction (q ≤ 0.05) are highlighted in blue. C) Foreground dN:dS ratio of positively selected codons (ω2) and statistical significance (P-value) from likelihood-ratio tests between positive-selection branch-site models and neutral branch-site models. Significant tests after FDR correction (q ≤ 0.05) are highlighted in orange. The red dashed lines in all plots represent neutral evolution (ω =1).

### Positive gene selection at the pathway level

We then visualized the interactions of the positive gene selection at a pathway level using IPA signaling pathways. Insulin signaling (Fig S1, Table S2), mTOR signaling (Fig S2, Table S3), NF-κB signaling (Fig S3, Table S4) and SIRT signaling (Fig S4, Table S5) were found to be positively selected, especially those related to glucose metabolism and inflammation. Genes particularly upstream from lipid metabolism, cell growth and proliferation and apoptosis functions were positively selected in the cetacean lineage. Little differences in selection were found for p53 signaling (Fig S5, Table S6) and leptin signaling (Fig S6, Table S7).

### Prediction of functional effects

Using the BEB method in PAML, a total of 1936 codons among the 133 genes positively selected in the cetacean lineage (mean 14.78 codons per gene) were under positive selection. The predicted functional effects of the amino-acid substitutions in cetacea were predominantly deleterious (median provean score: -2.9) and only 8.5 % of substitutions had a positive score (Fig 3).

**Figure 3.**
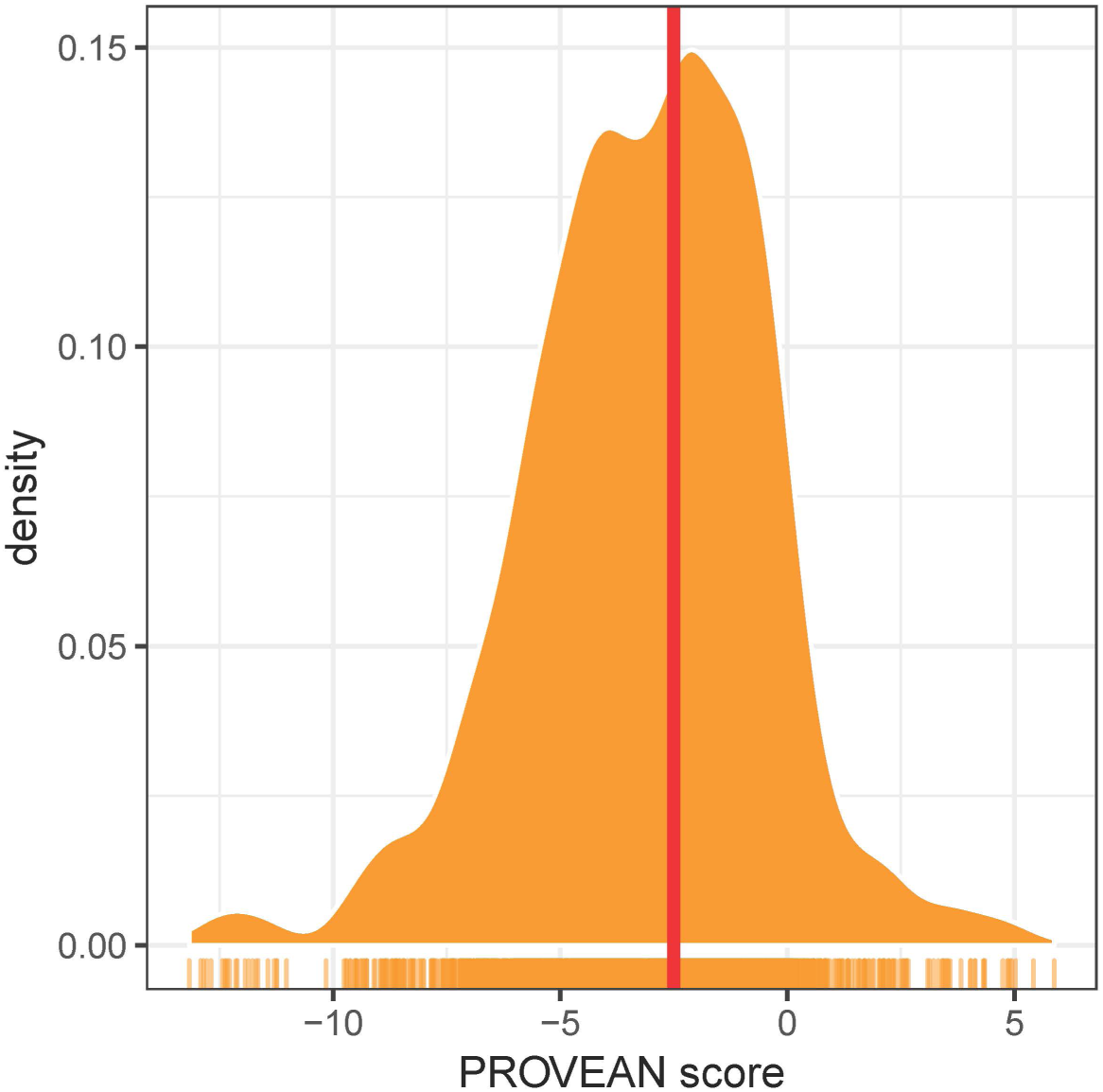
Distribution of *PROVEAN* scores of 1936 amino-acid substitutions under positive selection in cetacea. The red line indicates the standard threshold (−2.5) below which the effect of a substitution is considered deleterious.

These deleterious effects were likely to impact the way biological processes function. Unsurprisingly, glucose metabolism is expected to be altered with changes in insulin signaling (Fig 4), SIRT3 signaling with downstream regulation of insulin sensitivity, PPARA signaling with downstream regulation of gluconeogenesis and oxidation of fatty acids (Fig 5). In addition, we expect changes in upstream signaling of inflammation, hypoxia and cell survival via SIRT6, RB1, NFKB and HIF1a (Fig 5&6). Importantly, both the mTOR complexes and its upstream and downstream genes were estimated to be changed (Fig 7). The changes would have effects on nutrient sensing and protein synthesis as well as key biological decisions about energetic investment such as shift to glycolysis and de novo lipid synthesis.

**Figure 4.**
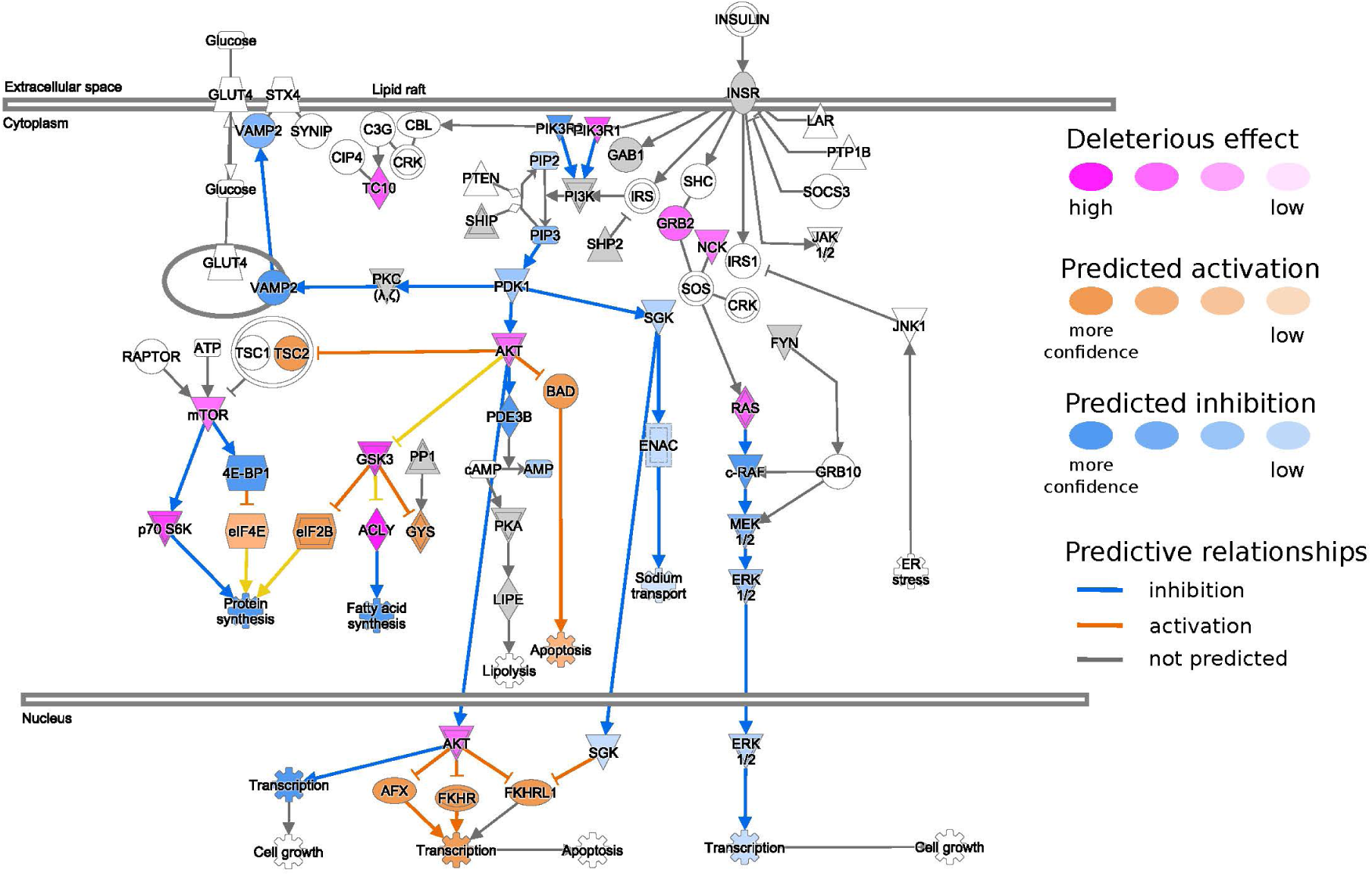
The insulin signaling pathway obtained from the Ingenuity Pathway Analysis (IPA) program. Genes with a damaging amino acid substitution are colored in magenta. Possible downstream damaging effects of these genes were visualized using the Molecule Activity Predictor (MAP) tool in IPA (see prediction legend).

**Figure 5.**
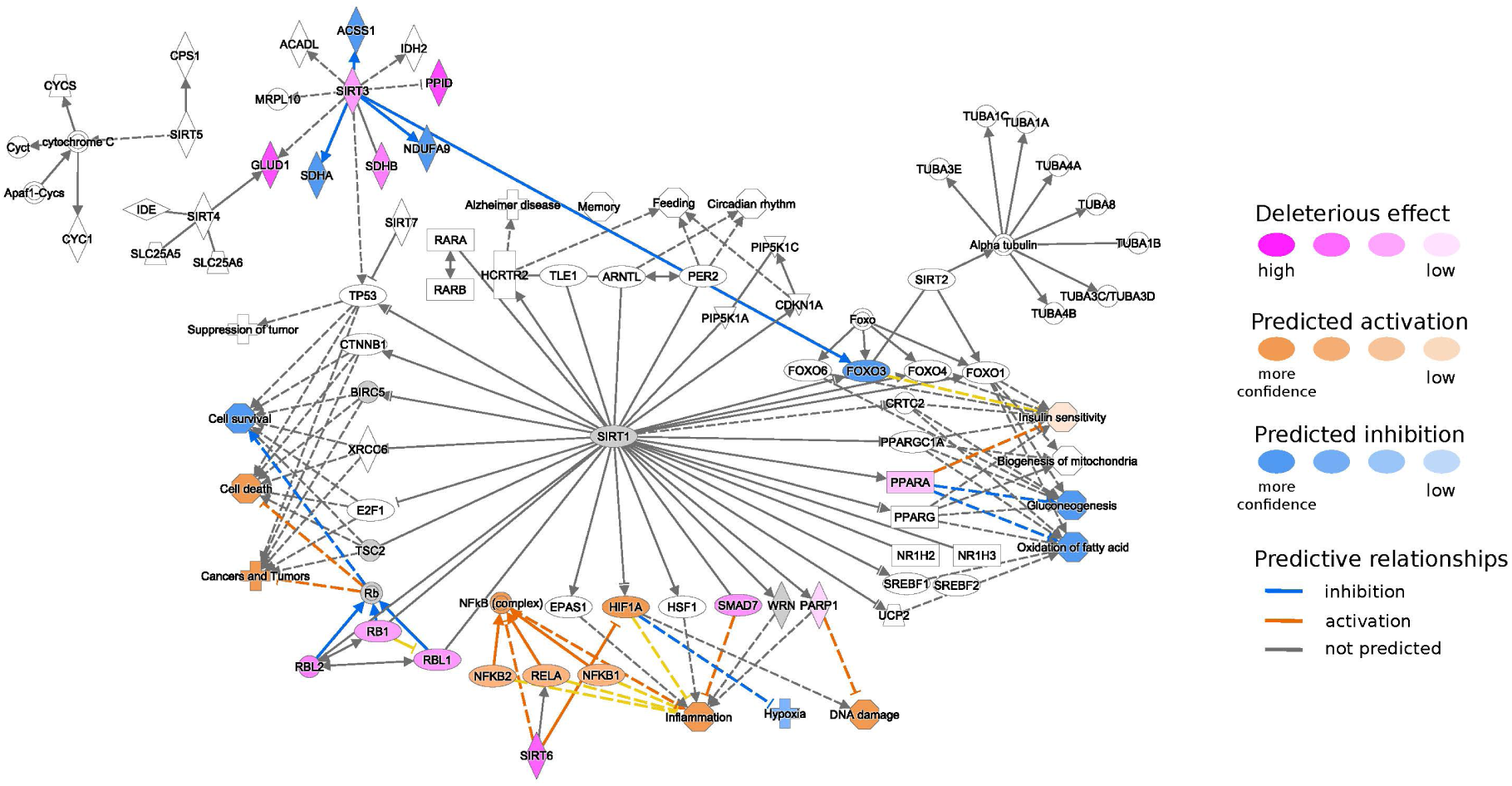
The SIRT signaling pathway obtained from the Ingenuity Pathway Analysis (IPA) program. Genes with a damaging amino acid substitution are colored in magenta. Possible downstream damaging effects of these genes were visualized using the Molecule Activity Predictor (MAP) tool in IPA (see prediction legend).

**Figure 6.**
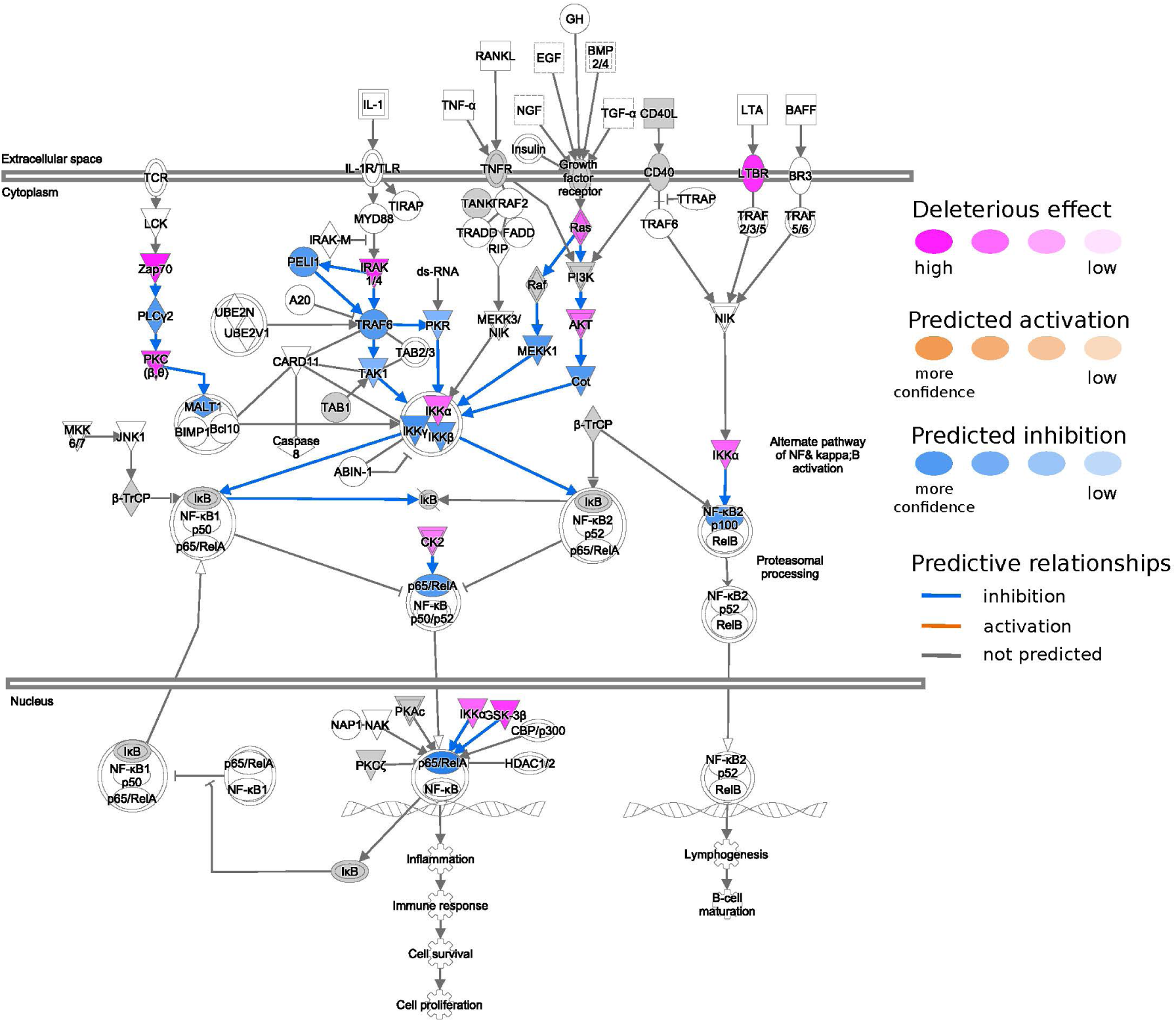
The NF-κB signaling pathway obtained for Ingenuity Pathway Analysis (IPA) program. Genes with a damaging amino acid substitution are colored in magenta. Possible downstream damaging effects of these genes were visualized using the Molecule Activity Predictor (MAP) tool in IPA (see prediction legend).

**Figure 7.**
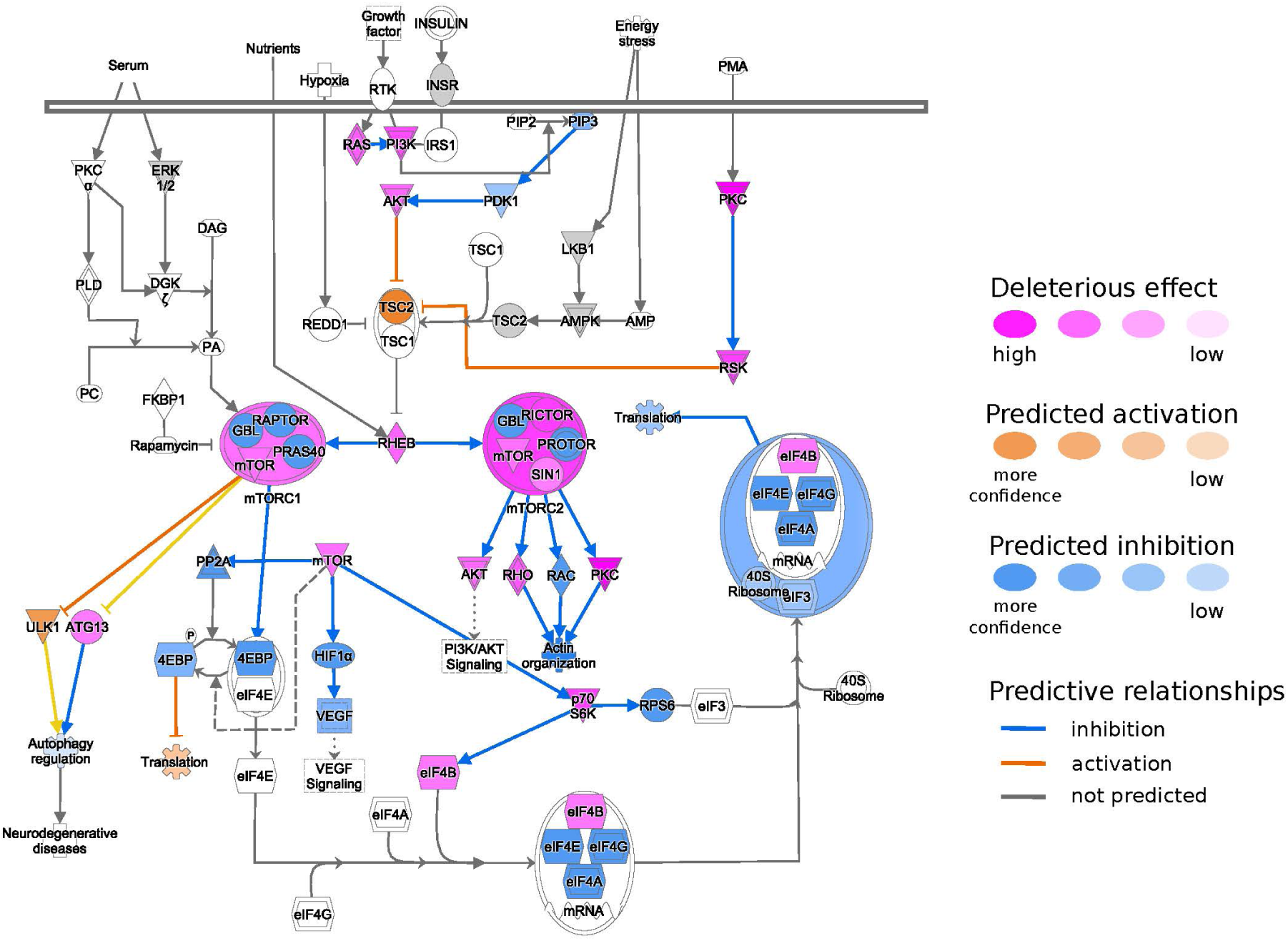
The mTOR signaling pathway obtained for Ingenuity Pathway Analysis (IPA) program. Genes with a damaging amino acid substitution are colored in magenta. Possible downstream damaging effects of these genes were visualized using the Molecule Activity Predictor (MAP) tool in IPA (see prediction legend).

## Discussion

Here we took a targeted approach to identify positive selected genes in cetacean nutrient sensing pathways. This allowed us to better understand how combined changes might be focused on particular functions within these pathways and what the consequences of these changes might be for the way by which energy metabolism in cetaceans may differ from their terrestrial counterparts. These pathways have signaling cascades in common with hormones such as insulin and are linked with the release of hormones from adipose tissue (e.g. leptin). Our results indicate that genes involved in the insulin signaling pathway, the mTOR, SIRT and NF-κB signaling pathway were significantly positively selected. These pathways have profound effects on metabolism and the maintenance of energy reserves. In terrestrial mammals, adipose tissue mass is related to metabolic fitness and expansion can lead to inflammation triggering metabolic disfunctions and diseases (e.g. obesity, metabolic syndrome, insulin resistance). The positive selection of genes related to glucose metabolism and inflammation suggest that these genes may have been positively selected to adapt to a glucose-poor diet and that fat deposits signaling may not be as limited by inflammation, metabolic dysfunctions (e.g. insulin resistance) and reproduction. Understanding these adaptations can help us manage conservation threats that perturb the environmental nutrient levels of cetaceans (National Academies of Sciences, Engineering, 2017).

Cetaceans have a diet with a high fat and protein content and is almost devoid of carbohydrates (Wells et al., 2013). Hence pathways regulating carbohydrate and glucose metabolism would have been under selective pressure as these species underwent a shift in substrate utilization. Genes related to the control of food intake, glycerol uptake and glucose metabolism were found to be under positive selective pressure in dolphins (McGowen, Grossman, & Wildman, 2012). A high glucose transport may be needed via erythrocytes to deliver glucose to specific brain regions under normal or physiological stress conditions (e.g. hypoxia while diving) (Craik, Young, & Cheeseman, 1998). We found positive selections in the insulin signaling pathway which are consistent with this hypothesis. In fasted Northern elephant seals components of the insulin signaling pathway were reduced including glucose transport (GLUT4), phosphatidylinositol 3-kinase (PI3K) and phosphorylated insulin receptor substrate 1 (IRS1) (Viscarra, Vázquez-Medina, Crocker, & Ortiz, 2011). In our study, both PI3K and IRS1 genes were estimated to be under positive selection but not GLUT4. Elephant seals are also insulin resistant (Viscarra et al., 2011) and hence may share common evolutionary selection of those mechanisms with cetaceans. We indeed identified deleterious changes to the insulin signaling pathway, including the Akt protein signaling kinase (Akt) and PI3K. Damage to this pathway in various tissues has been linked to insulin resistance (Huang, Liu, Guo, & Su, 2018). The three Akt isoforms have differential physiological functions but loss of function in one isoform is compensated by another. Both Akt2^-/-^ and Akt3^-/-^ mice exhibit severe glucose and insulin resistance (Dummler et al., 2006). Interestingly the Akt phosphorylation at Ser473, which is required for its full activation, is accomplished by mTORC2 (Oh & Jacinto, 2011; Sarbassov, Guertin, Ali, & Sabatini, 2005). Both mTORC1 and mTORC2 were positively selected in the cetacean lineage to a point where we cannot expect them to function in the same way as in terrestrial mammals. In addition to PI3K/AKT signaling, PPARα was also positively selected. Activation of the PPARα isoform leads to improved lipid and carbohydrate profile and to reduced inflammation (Moller & Berger, 2003). It’s anti-inflammatory properties have been linked to the suppression of NF-κB (Fuentes, Guzmán-Jofre, Moore-Carrasco, & Palomo, 2013). However, PPARα is predominantly involved in cellular uptake, activation and β-oxidation of fatty acids (Moller & Berger, 2003). PPARα^-/-^ mice exposed to long-term high fat diet remained normoglycemic and normoinsulinemic despite having high adiposity while the wild type developed hyperinsulemia. In addition, glucose and insulin tolerance test indicated that high-fat-fed wild type developed insulin resistance over time while the PPARα^-/-^ remained unchanged (Guerre-Millo et al., 2001). Hence in absence of PPARα, the increase in adiposity as a result of a high fat diet does not lead to insulin resistance. Interestingly, when dolphins were fed a big meal of fish, they do show signs of insulin resistant (Venn-Watson et al., 2011). However, when just fed dextrose and water they showed an insulin-deficient response (S. Venn-Watson et al., 2013). This response to glucose is similar to the GTT test of the PPARα^-/-^ mice. Hence key genes such as PPARα, AKT and PI3K in the insulin signaling pathway may be positively selected as an evolutionary driver for insulin resistance and to switch between type 2 and type 1 diabetes like states (S. Venn-Watson, 2014).

Maintaining adiposity can have negative consequences for survival in terrestrial mammals. We know that a large volume of adipose tissue triggers inflammatory responses in a range of species and can lead to metabolic dysfunctions at a physiological level (e.g. insulin resistance) (Mantovani, Sozzani, Locati, Allavena, & Sica, 2002). NF-κB is involved in the molecular signaling of hypoxia during adipose tissue expansion and triggers these inflammatory responses (Ye, Gao, Yin, & He, 2007). As the inflammatory function of NF-κB is linked to fat mass, the high level of adiposity in cetaceans would lead to chronic inflammation. The thickened blubber of cetaceans is a result of the secondary adaptation to life in water and hence selective pressure in this pathway may be a way to reduce intrinsic tissue inflammation. Here we found that key genes in the NF-κB signaling pathway were positively selected inhibitor of nuclear factor kappa B kinase subunit beta (IKKβ) and IKKα. Mice that have the inflammatory pathway of NF-κB disabled (IKKβ knockout) are more insulin sensitive and are partially protected from high fat diet induced glucose intolerance and hyperinsulinemia (Arkan et al., 2005). In addition, the gene regulation of receptor interacting serine/threonine kinase 1 (RIPK1) was also positively selected and its associated amino acid sequence changed drastically compared to the outgroups. RIPK1 has a downstream effect on IKKα and IKKβ, and hence may influence the signaling in this pathway. The IKK complex has a NF-κB independent role in the protection of cells from RIPK–dependent death downstream from the tumor necrosis factor rector (TNFR1) (Dondelinger et al., 2015). Cetaceans do have a fully functioning immune and endocrine responses and these are highly dependent on the environment (e.g. pathogens, pollution and noise) (Fair & Becker, 2000; Fair et al., 2017). For example transcripts encoding pro-inflammatory cytokines were significantly lower in managed-care dolphins compared to free-ranging dolphins (Fair et al., 2017). A greater understanding of tissue specific inflammatory responses is needed and may provide valuable insights into how inflammatory responses are regulated in regards to the high adiposity in these healthy but “obese” mammals, during periods of diving and environmental fluctuation.

Finally, most components in the mTOR pathway were positively selected including both of its complexes (mTORC1 and mTORC2). mTORC1 regulates processes related to growth and differentiation while mTORC2 plays a regulatory role in the insulin cascade (Lamming et al., 2012). mTOR is primarily involved in immune response and sensing nutrient availability. As elevated mTOR leads to increased hepatic gluconeogenesis and reduced glucose uptake by muscles, it is maybe not surprising that several components in the mTOR pathway are significantly changed in cetaceans. Especially as mTOR is involved in nutrient sensing. As dolphins are able to switch between type 2 and type 1 diabetes like states based on their meal content (S. Venn-Watson, 2014), specific components both up- and down-stream from mTOR may be positively selected to facilitate such a response. In rats, a ketogenic diet (low in carbohydrates) are able to reduce mTOR expression and likely via AKT (McDaniel, Rensing, Thio, Yamada, & Wong, 2011). Rodents fed on a keto diet also exhibit lower insulin levels, which likely induce a decreased mTOR signaling (Thio, Erbayat-Altay, Rensing, & Yamada, 2006). Cetaceans have a diet similar to ketogenic diet with a high fat and protein content and almost devoid of carbohydrates (Wells et al., 2013). Hence, to optimise the uptake of the limited available glucose in the diet, aspects in the insulin/mTOR pathway may be altered to create insulin resistance.

## Conclusion

Taken together, our findings provide novel insights into the role of the insulin, mTOR and NF-κB signaling pathways in the adaptation of cetaceans to an aquatic life. They point to adaptations likely to reduce the health consequences of adiposity. These results mean that condition measures based on adiposity must be used with caution. Indeed, lower bounds of adiposity are influenced by thermoregulatory requirements and upper bounds of adiposity will not be influenced by inflammatory response in the same way as it is in terrestrial mammals.

Further work is needed to unravel the complex signaling mechanisms of adipose tissue in cetacean energy metabolism and to determine the effects of these signaling molecules on whole body functioning including appetite regulation, energy balance, and inflammatory responses. Understanding these adaptations can help us manage conservation threats that perturb the environmental nutrient levels of cetaceans (National Academies of Sciences, Engineering, 2017).

## Authors’ Contributions

DL, JS and DD designed the study. DD wrote the manuscript and performed the pathway level analyses and interpretation with input from DL. MW performed the gene-level analyses with input from AD and JS. DD, MW and DL interpreted the data. All authors read and commented on the manuscript.

## Acknowledgements

This work was funded by US ONR grant award number N000141512377. The authors acknowledge the support of the Maxwell computer cluster funded by the University of Aberdeen.

## Data accessibility

Data will become openly available after uploading on Dryad.

## Supplementary Figures

**Figure S1.**
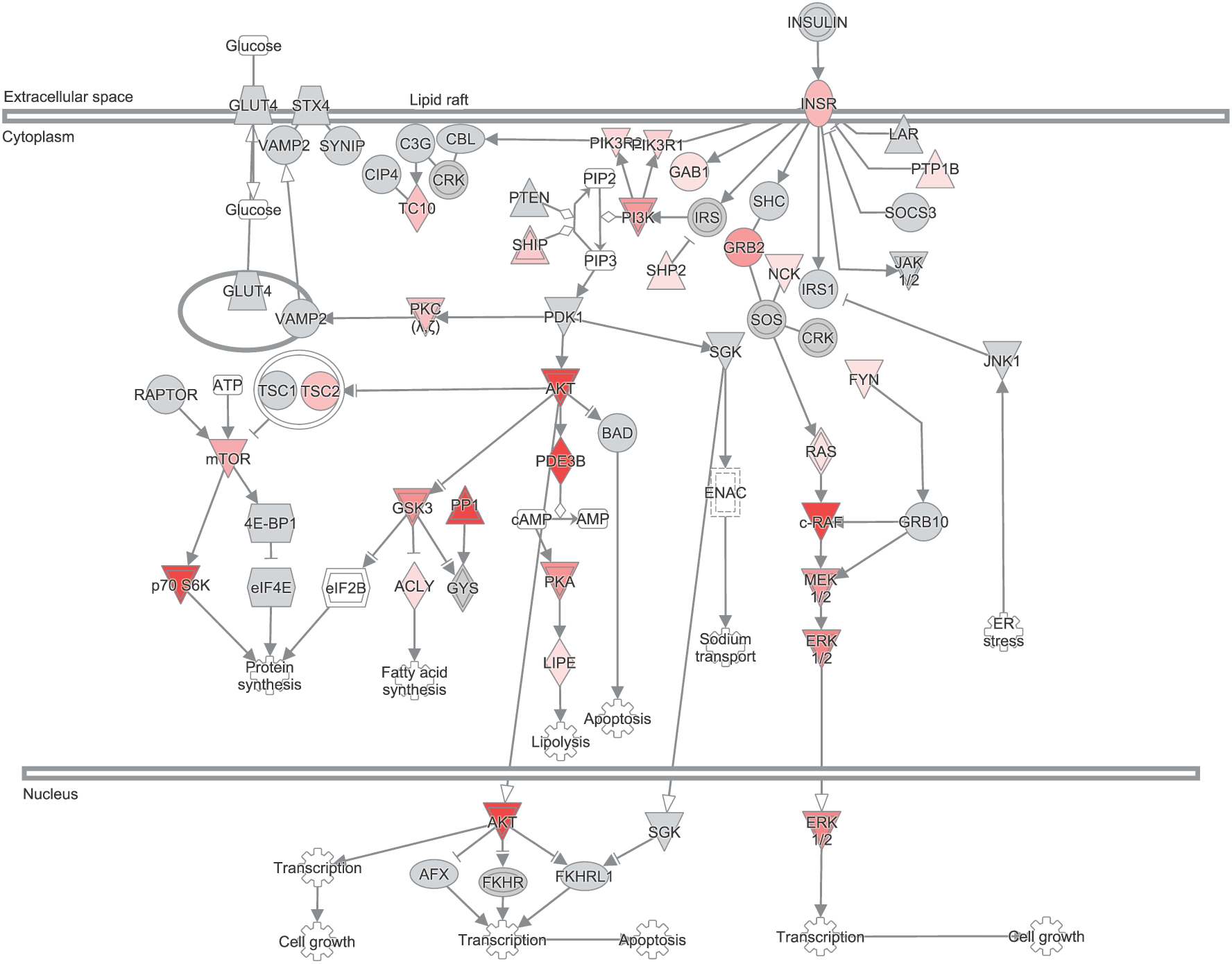
The insulin signaling pathway obtained from Ingenuity Pathway Analysis (IPA) program. Genes in the pathway are colored according to their corresponding 2ΔL value identified by the branch-site model. Intensity of the color is related to the strength of the positive gene selection. Uncolored genes represent those genes with an adjusted p-value > 0.05.

**Figure S2.**
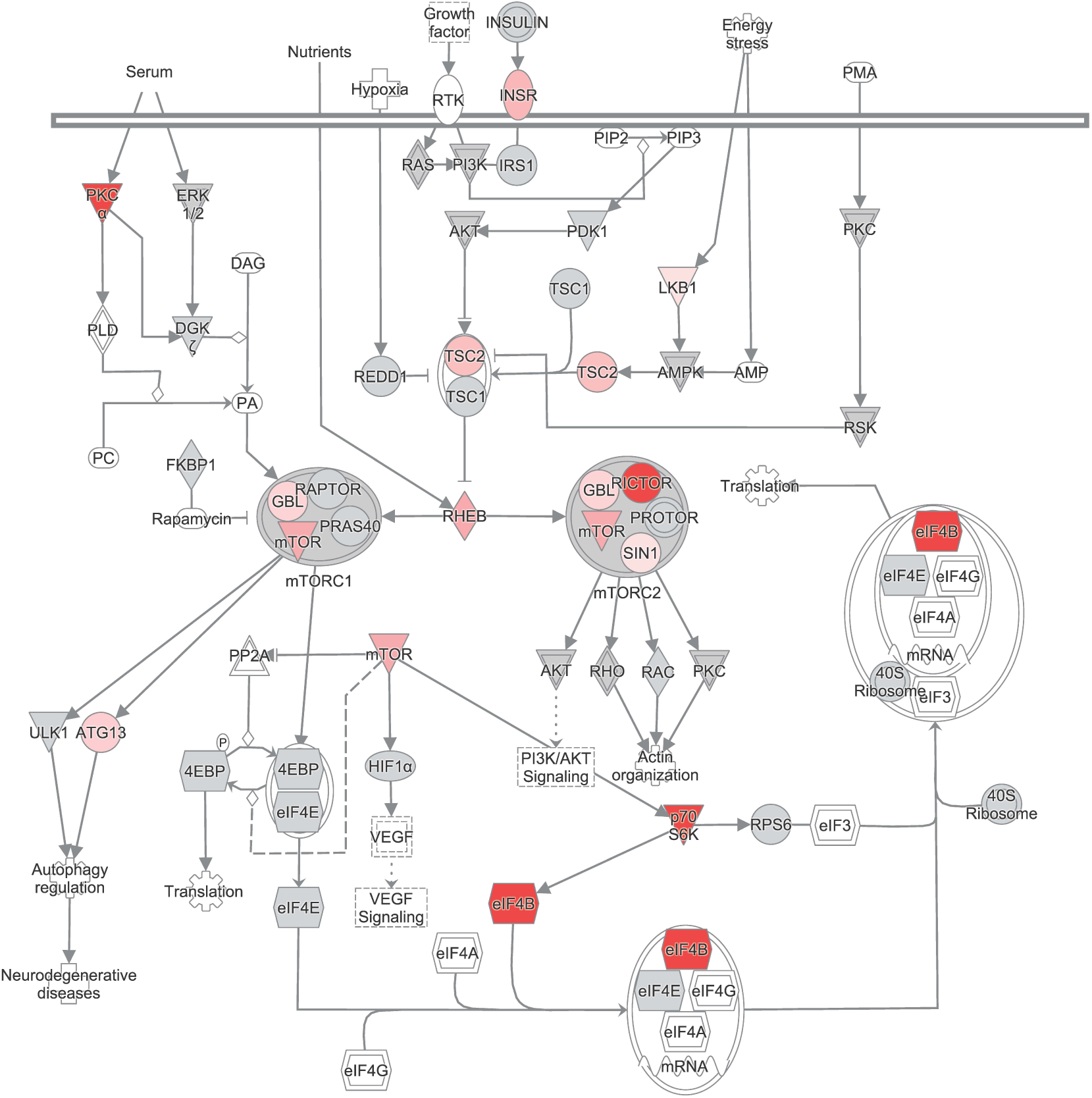
The mTOR signaling pathway obtained for Ingenuity Pathway Analysis (IPA) program. Genes in the pathway are colored according to their corresponding 2ΔL value identified by the branch-site model. Intensity of the color is related to the strength of the positive gene selection. Uncolored genes represent those genes with an adjusted p-value > 0.05.

**Figure S3.**
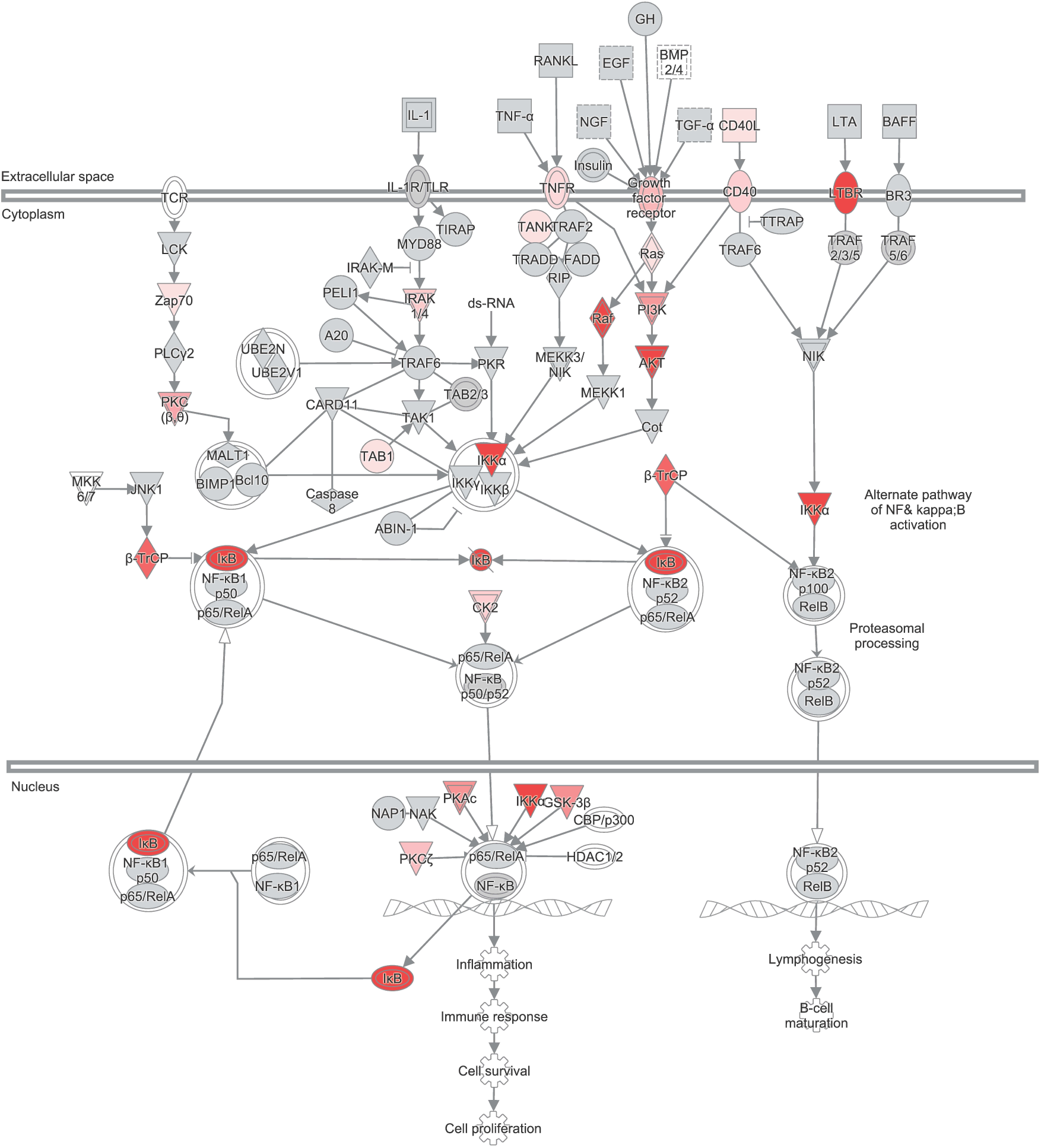
The NF-κB signaling pathway obtained for Ingenuity Pathway Analysis (IPA) program. Genes in the pathway are colored according to their corresponding 2ΔL value identified by the branch-site model. Intensity of the color is related to the strength of the positive gene selection.

**Figure S4.**
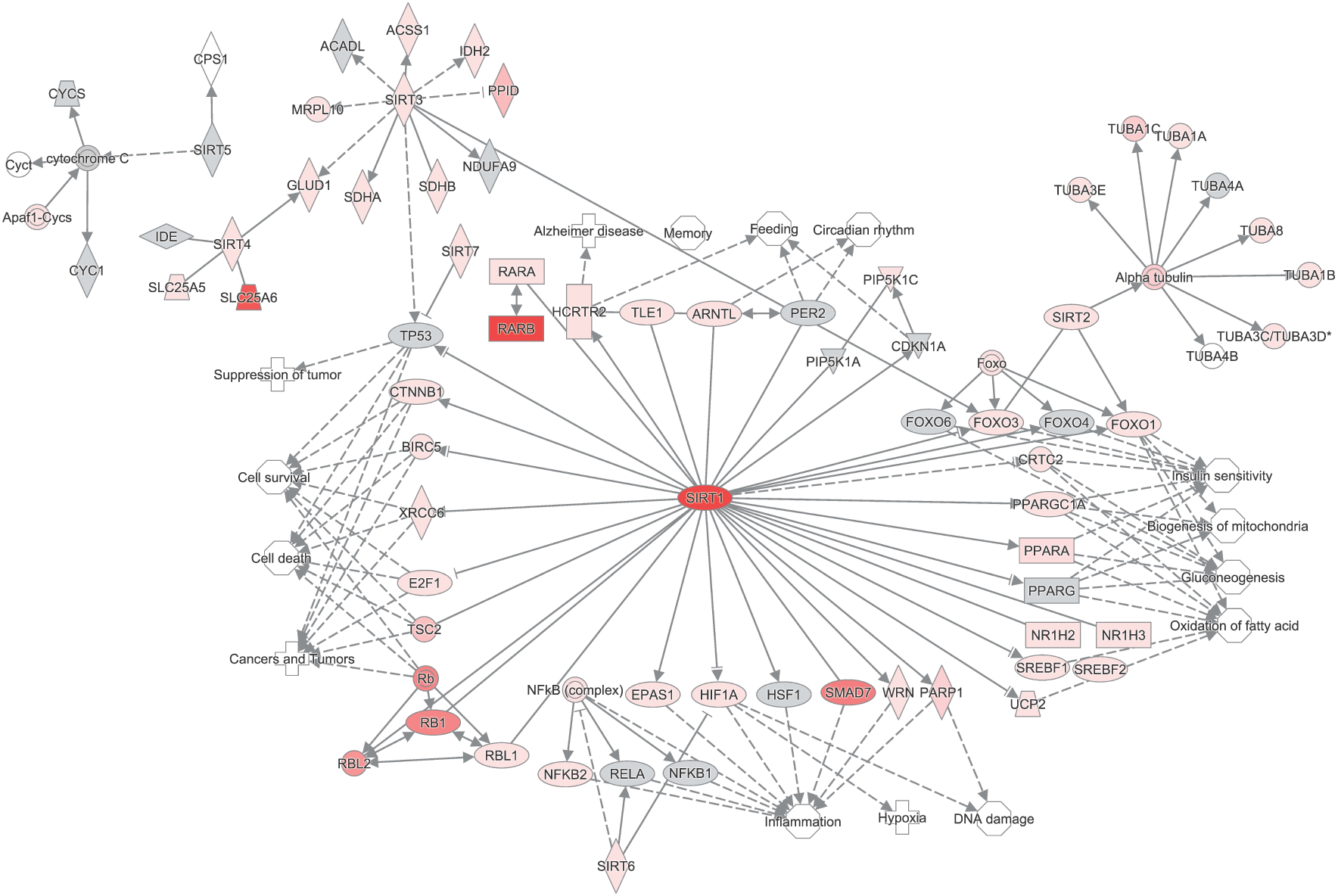
The SIRT signaling pathway created in the Ingenuity Pathway Analysis (IPA) program. Genes in the pathway are colored according to their corresponding 2ΔL value identified by the branch-site model. Intensity of the color is related to the strength of the positive gene selection.

**Figure S5.**
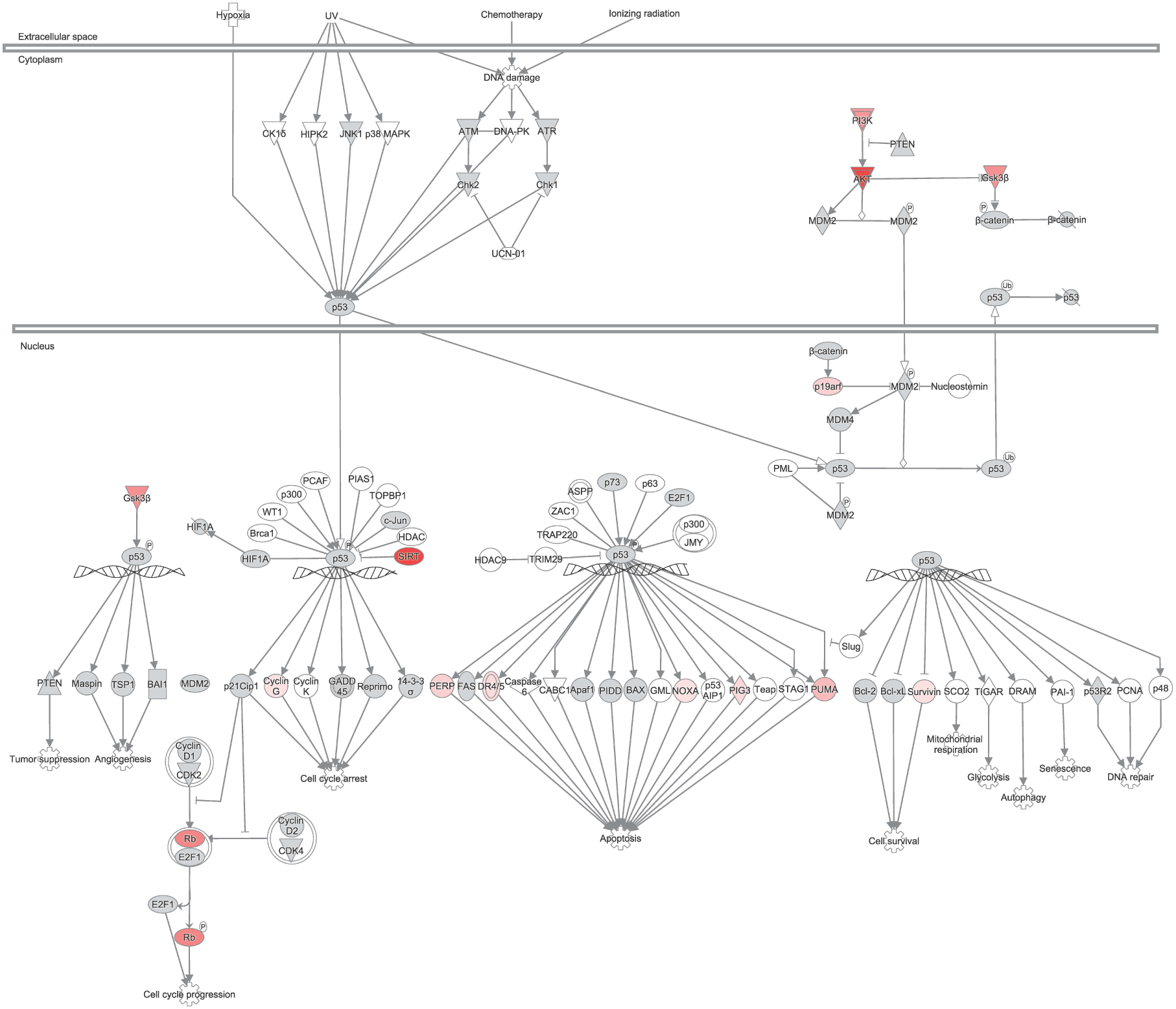
The p53 signaling pathway obtained for Ingenuity Pathway Analysis (IPA) program. Genes in the pathway are colored according to their corresponding 2ΔL value identified by the branch-site model. Intensity of the color is related to the strength of the positive gene selection.

**Figure S6.**
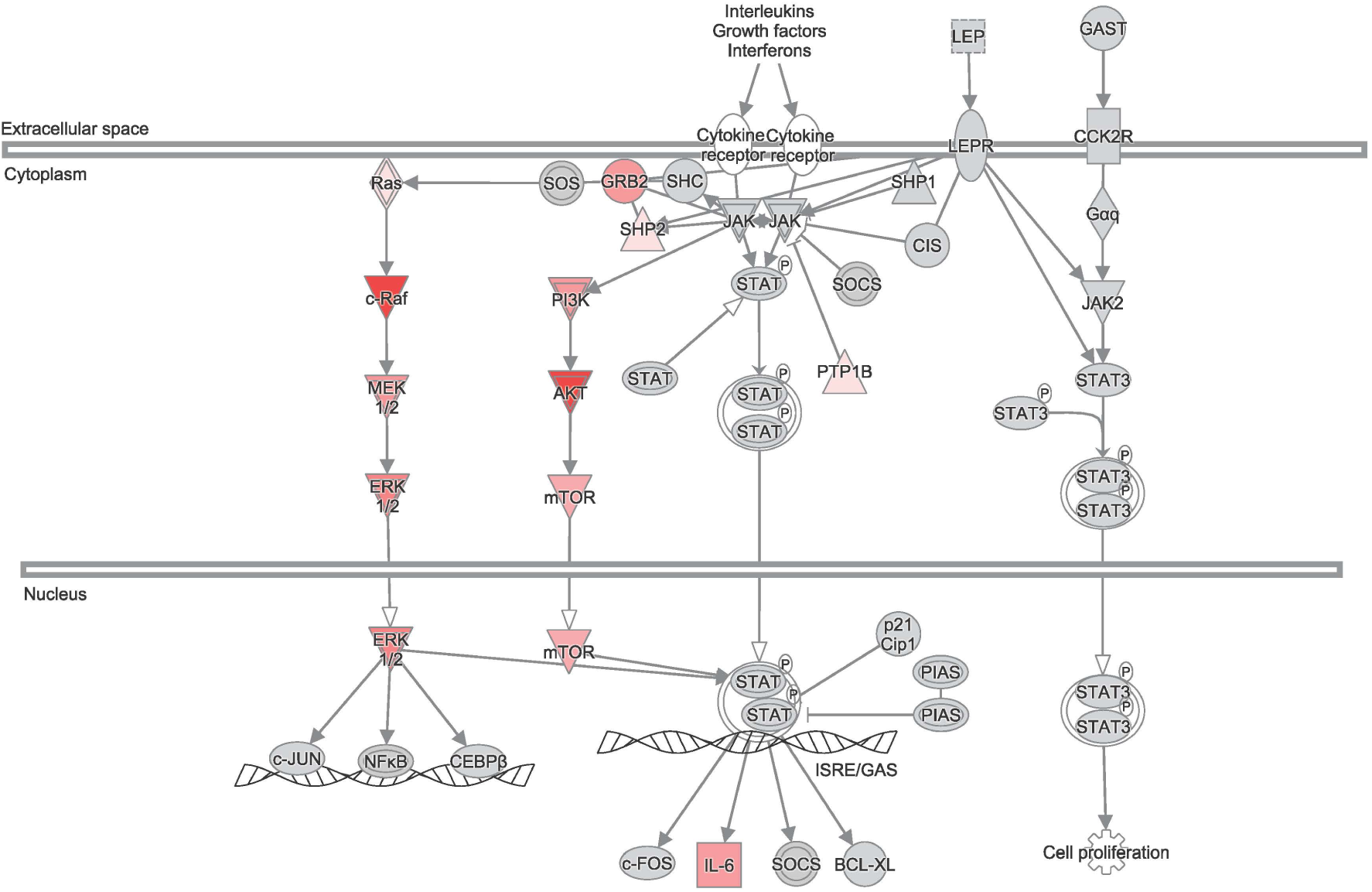
The leptin signaling pathway obtained for Ingenuity Pathway Analysis (IPA) program. Genes in the pathway are colored according to their corresponding 2ΔL value identified by the branch-site model. Intensity of the color is related to the strength of the positive gene selection.

## Supplementary Tables

**Table S1.**
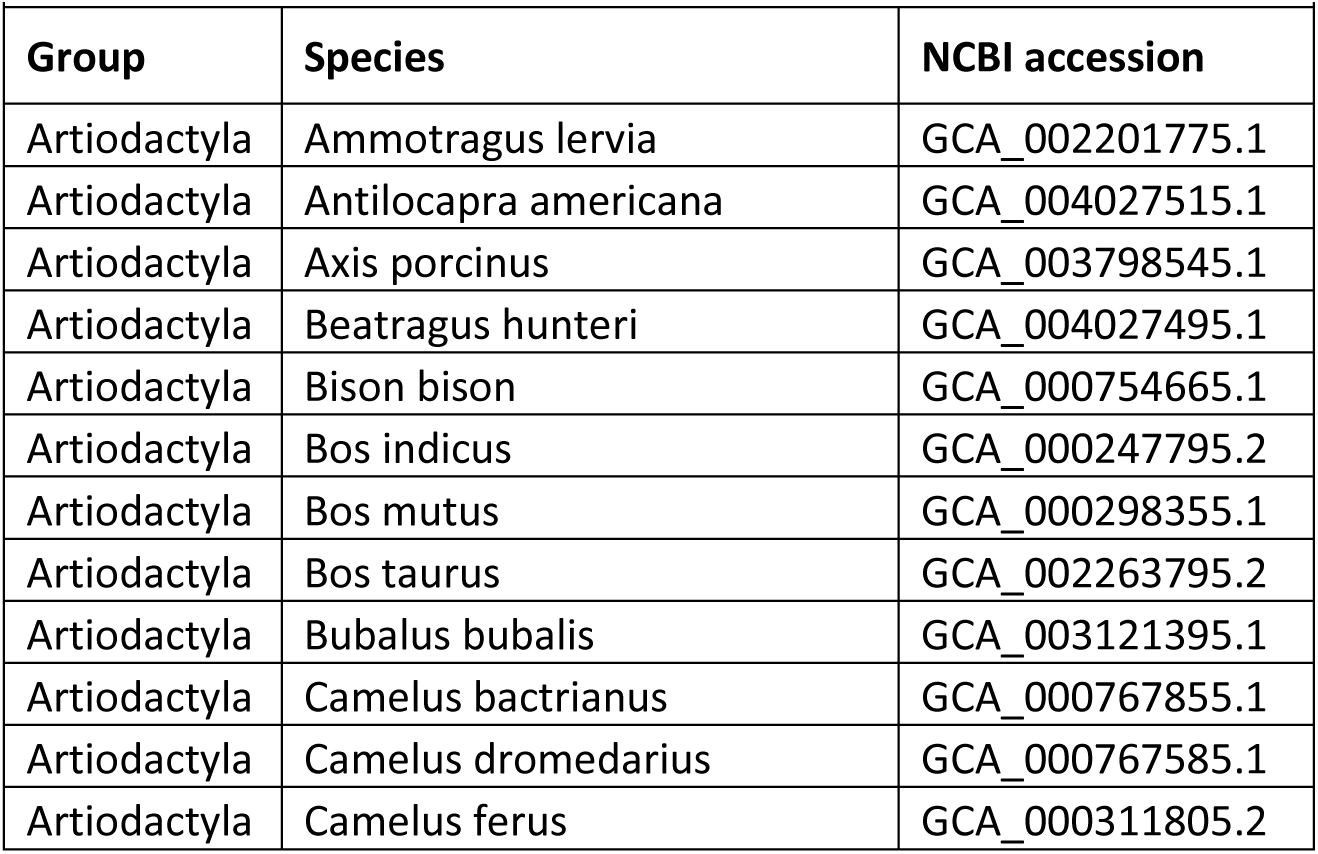
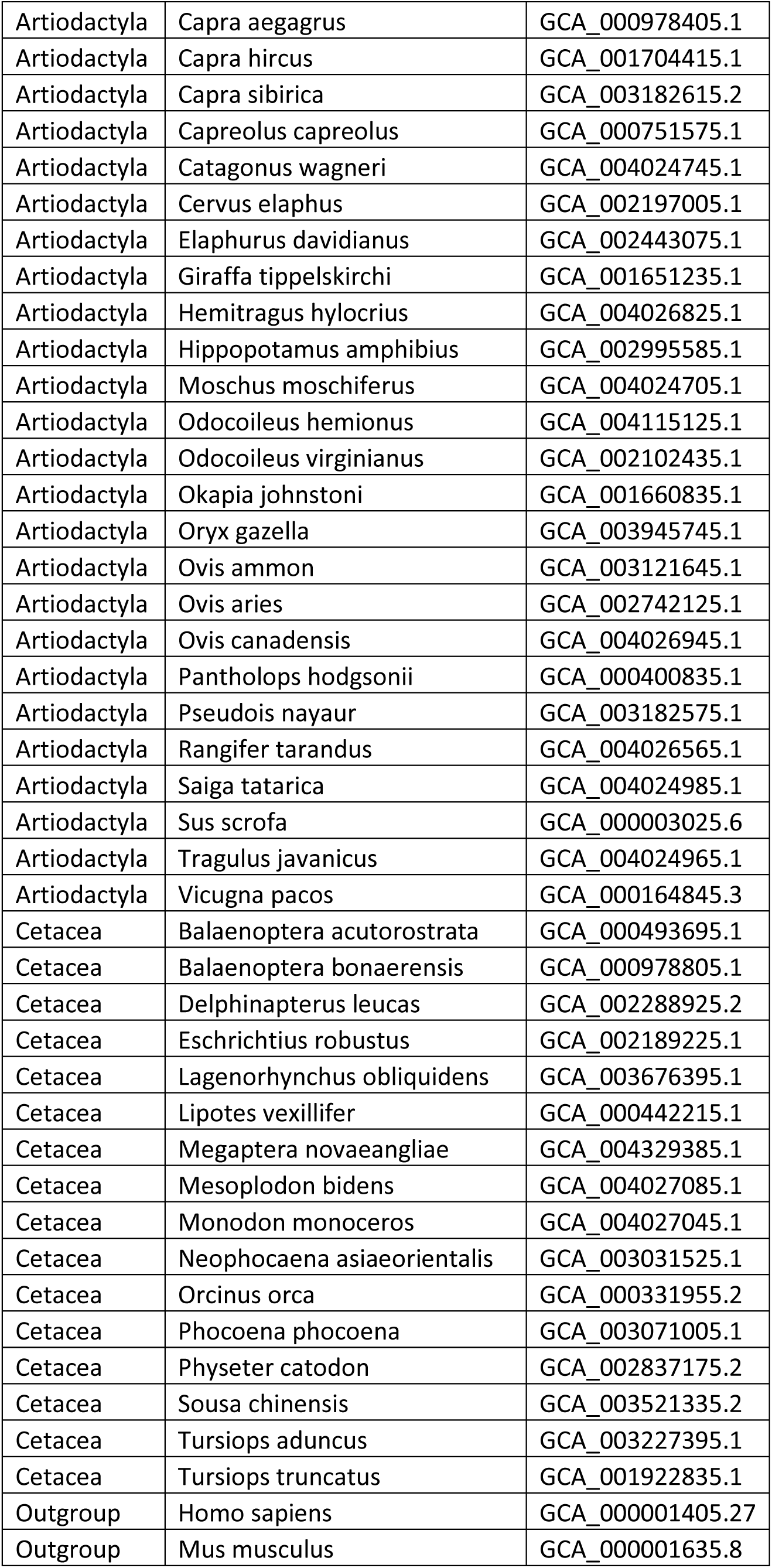
The genome assemblies of human, mouse, 16 cetacean species and 37 artiodactyl species downloaded from NCBI.

**Table S2.** Insulin signaling pathway overlaid with the likelihood-ratio test statistics (2ΔL) between the positive and neutral branch-models as a measure to visualize a pathway effect.

| **Symbol** | **$2\Delta L$** | ***p*-value** |
| --- | --- | --- |
| 4E-BP1 | 0.000 | 1.000 |
| ACLY | 17.800 | <0.001 |
| AFX | 0.000 | 1.000 |
| AKT |  |  |
| AMP |  |  |
| Apoptosis |  |  |
| ATP |  |  |
| BAD | 1.359 | 0.566 |
| c-RAF | 105.387 | <0.001 |
| C3G | 0.156 | 1.000 |
| cAMP |  |  |
| CBL | 0.000 | 1.000 |
| Cell growth |  |  |
| CIP4 | 0.003 | 1.000 |
| CRK |  |  |
| eIF2B |  |  |
| eIF4E | 0.333 | 1.000 |
| ENAC |  |  |
| ER stress |  |  |
| ERK1/2 |  |  |
| Fatty acid synthesis |  |  |
| FKHR |  |  |
| FKHRL1 | 0.000 | 1.000 |
| FYN | 8.207 | 0.016 |
| GAB1 | 11.223 | 0.004 |
| Glucose |  |  |
| GLUT4 | 0.188 | 1.000 |
| GRB10 | 3.968 | 0.139 |
| GRB2 | 50.205 | <0.001 |
| GSK3 |  |  |
| GYS |  |  |
| INSR | 34.960 | <0.001 |
| INSULIN |  |  |
| IRS |  |  |
| IRS1 | 0.001 | 1.000 |
| JAK1/2 |  |  |
| JNK1 | 0.001 | 1.000 |
| LAR | 0.000 | 1.000 |
| LIPE | 16.254 | <0.001 |
| Lipolysis |  |  |
| MEK1/2 |  |  |
| mTOR | 41.010 | <0.001 |
| NCK | 9.400 | 0.009 |
| p70 S6K |  |  |
| PDE3B | 105.839 | <0.001 |
| PDK1 | 0.000 | 1.000 |
| PI3K |  |  |
| PIK3R1 | 22.707 | <0.001 |
| PIK3R2 | 20.576 | <0.001 |
| PIP2 |  |  |
| PIP3 |  |  |
| PKA |  |  |
| PKC($\lambda, \zeta$) | | |
| PP1 |  |  |
| Protein synthesis |  |  |
| PTEN | 0.019 | 1.000 |
| PTP1B | 14.851 | 0.001 |
| RAPTOR | 0.318 | 1.000 |
| RAS |  |  |
| SGK | 0.110 | 1.000 |
| SHC | 0.000 | 1.000 |
| SHIP |  |  |
| SHP2 | 9.205 | 0.009 |
| SOCS3 | 0.000 | 1.000 |
| Sodium transport |  |  |
| SOS |  |  |
| STX4 | 0.000 | 1.000 |
| SYNIP | 0.000 | 1.000 |
| TC10 | 31.276 | <0.001 |
| Transcription |  |  |
| TSC1 | 4.121 | 0.129 |
| Tsc1-Tsc2 |  |  |
| TSC2 | 31.417 | <0.001 |
| VAMP2 | 3.423 | 0.185 |

**Table S3.** mTOR signaling pathway overlaid with the likelihood-ratio test statistics (2ΔL) between the positive and neutral branch-models as a measure to visualize a pathway effect.

| Symbol | $2\Delta L$ | $p$-value |
| --- | --- | --- |
| 40S Ribosome-eIF3-mRNA-eIF4A-eIF4B-eIF4E-eIF4G |  |  |
| 40SRibosome |  |  |
| 4EBP | 0.000 | 1.000 |
| 4EBP-eIF4E |  |  |
| Actin organization |  |  |
| AKT |  |  |
| AMP |  |  |
| AMPK |  |  |
| ATG13 | 24.238 | <0.001 |
| Autophagy regulation |  |  |
| DAG |  |  |
| DGKζ | 4.209 | 0.124 |
| eIF3 |  |  |
| eIF4A |  |  |
| eIF4A-eIF4B-eIF4E-eIF4G |  |  |
| eIF4B | 232.205 | <0.001 |
| eIF4E | 0.333 | 1.000 |
| eIF4G |  |  |
| ERK1/2 |  |  |
| FKBP1 | 0.010 | 1.000 |
| GBL | 20.882 | <0.001 |
| HIF1α | 0.000 | 1.000 |
| Hypoxia |  |  |
| INSR | 34.960 | <0.001 |
| INSULIN |  |  |
| IRS1 | 0.001 | 1.000 |
| LKB1 | 11.489 | 0.003 |
| mTOR | 41.010 | <0.001 |
| mTORC1 |  |  |
| mTORC2 |  |  |
| Neurodegenerative diseases |  |  |
| p70S6K | 292.275 | <0.001 |
| PA |  |  |
| PC |  |  |
| PDK1 | 0.000 | 1.000 |
| PI3K |  |  |
| PIP2 |  |  |
| PIP3 |  |  |
| PKC |  |  |
| PKCα | 383.006 | <0.001 |
| PLD |  |  |
| PMA |  |  |
| PP2A |  |  |
| PRAS40 | 0.000 | 1.000 |
| PROTOR |  |  |
| RAC | 3.675 | 0.161 |
| Rapamycin |  |  |
| RAPTOR | 0.318 | 1.000 |
| RAS |  |  |
| REDD1 | 0.000 | 1.000 |
| RHEB | 42.036 | <0.001 |
| RHO |  |  |
| RICTOR | 472.142 | <0.001 |
| RPS6 | 1.495 | 0.531 |
| RSK |  |  |
| SIN1 | 10.430 | 0.005 |
| Translation |  |  |
| TSC1 | 4.121 | 0.129 |
| Tsc1-Tsc2 |  |  |
| TSC2 | 31.417 | <0.001 |
| ULK1 | 0.000 | 1.000 |
| VEGF |  |  |

**Table S4.** NF-κB signaling pathway overlaid with the likelihood-ratio test statistics (2ΔL) between the positive and neutral branch-models as a measure to visualize a pathway effect.

| Symbol | $2\Delta L$ | *p*-value |
| --- | --- | --- |
| β-TrCP | 77.820 | <0.001 |
| A20 | 0.000 | 1.000 |
| ABIN-1 | 4.063 | 0.132 |
| AKT |  |  |
| B-cell maturation |  |  |
| BAFF | 5.391 | 0.066 |
| Bcl10 | 0.000 | 1.000 |
| Bcl10-Card10-Malt1 |  |  |
| BIMP1 | 4.177 | 0.125 |
| BMP2/4 |  |  |
| BR3 | 5.818 | 0.053 |
| CARD11 | 1.116 | 0.639 |
| Caspase8 | 0.904 | 0.723 |
| CBP/p300 |  |  |
| CD40 | 24.403 | <0.001 |
| CD40L | 11.317 | 0.004 |
| Cell proliferation |  |  |
| Cell survival |  |  |
| Chuk-Ikbkb-Ikbkg |  |  |
| CK2 |  |  |
| Cot | 0.624 | 0.876 |
| EGF | 3.083 | 0.219 |
| FADD | 0.000 | 1.000 |
| GH | 0.000 | 1.000 |
| Growth factor receptor |  |  |
| GSK-3β | 55.087 | <0.001 |
| HDAC1/2 |  |  |
| IκB | | |
| IκB-NfκB1-RelA | | |
| IκB-NfκB2-RelA | | |
| IKKα | 129.661 | <0.001 |
| IKKβ | 0.014 | 1.000 |
| IKKγ | 2.158 | 0.368 |
| IL-1 |  |  |
| IL-1R/TLR |  |  |
| Immune response |  |  |
| Inflammation |  |  |
| Insulin |  |  |
| IRAK1/4 |  |  |
| IRAK-M | 0.000 | 1.000 |
| JNK1 | 0.001 | 1.000 |
| LCK | 0.000 | 1.000 |
| LTA | 1.123 | 0.639 |
| LTBR | 570.904 | <0.001 |
| Lymphogenesis |  |  |
| MALT1 | 0.000 | 1.000 |
| MEKK1 | 0.000 | 1.000 |
| MEKK3/NIK |  |  |
| MKK6/7 |  |  |
| MYD88 | 1.355 | 0.566 |
| NAK | 0.023 | 1.000 |
| NAP1 | 0.208 | 1.000 |
| NF-κB p50/p52 | | |
| NF-κB1 | 0.000 | 1.000 |
| NF-κB2 p100 | 0.000 | 1.000 |
| NfκB-RelA | | |
| NfκB1-RelA | | |
| NfκB2(p52)-RelB | | |
| NGF | 0.145 | 1.000 |
| NIK |  |  |
| p65/RelA | 0.000 | 1.000 |
| PELI1 | 0.000 | 1.000 |
| PI3K |  |  |
| PKAc |  |  |
| PKC($\beta$,$\theta$) | | |
| PKCζ | 31.135 | <0.001 |
| PKR | 0.000 | 1.000 |
| PLCγ2 | 0.000 | 1.000 |
| Raf |  |  |
| RANKL | 1.048 | 0.670 |
| Ras |  |  |
| RelB | 0.500 | 0.945 |
| RIP | 5.280 | 0.070 |
| TAB1 | 9.539 | 0.008 |
| TAB2/3 |  |  |
| TAK1 | 1.581 | 0.504 |
| TANK | 8.987 | 0.010 |
| TCR |  |  |
| TGF-α | 1.477 | 0.535 |
| TIRAP | 0.000 | 1.000 |
| TNF-α | 0.420 | 1.000 |
| TNFR |  |  |
| TRADD | 2.123 | 0.375 |
| TRAF2/3/5 |  |  |
| TRAF5/6 |  |  |
| TRAF2 | 0.000 | 1.000 |
| TRAF6 | 1.795 | 0.444 |
| TTRAP | 0.000 | 1.000 |
| UBE2N | 0.011 | 1.000 |
| Ube2n-Ube2v1 |  |  |
| UBE2V1 | 0.000 | 1.000 |
| Zap70 | 11.445 | 0.003 |

**Table S5.**
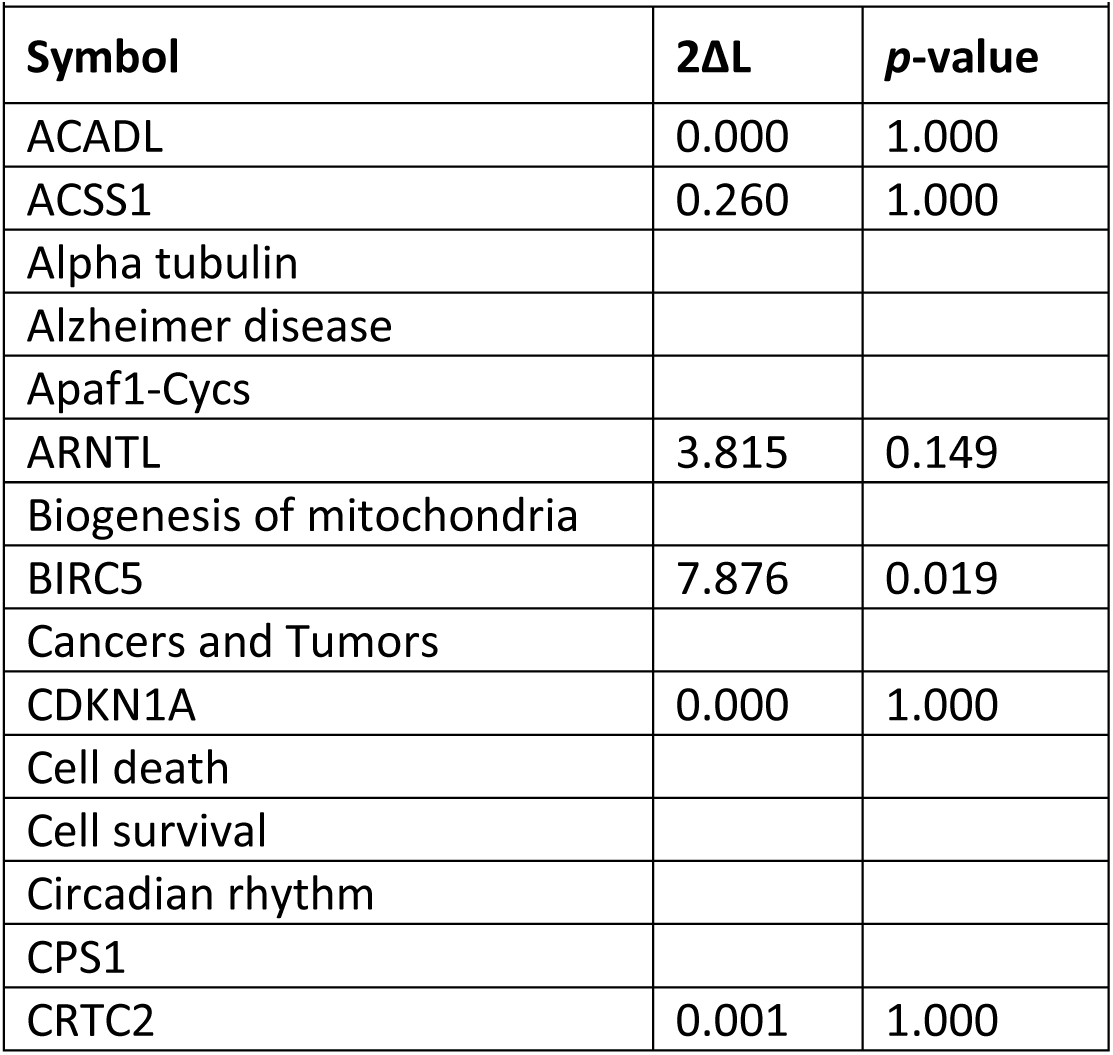
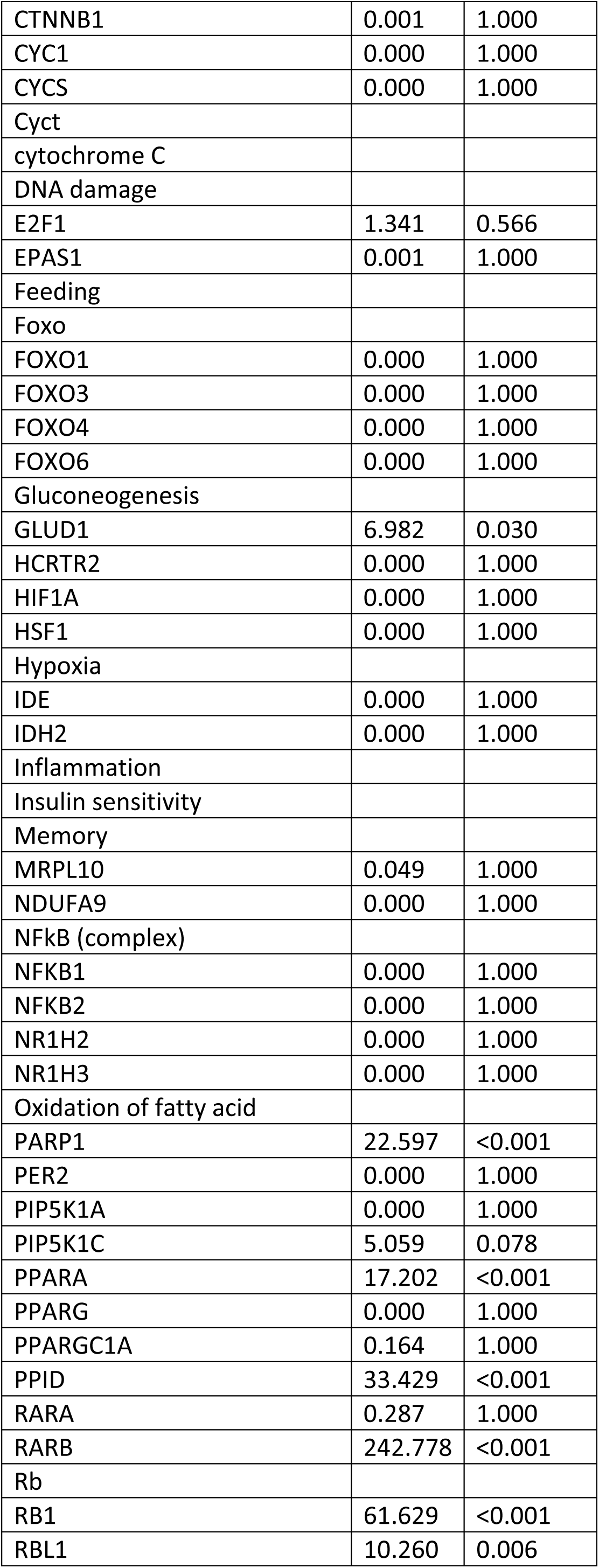
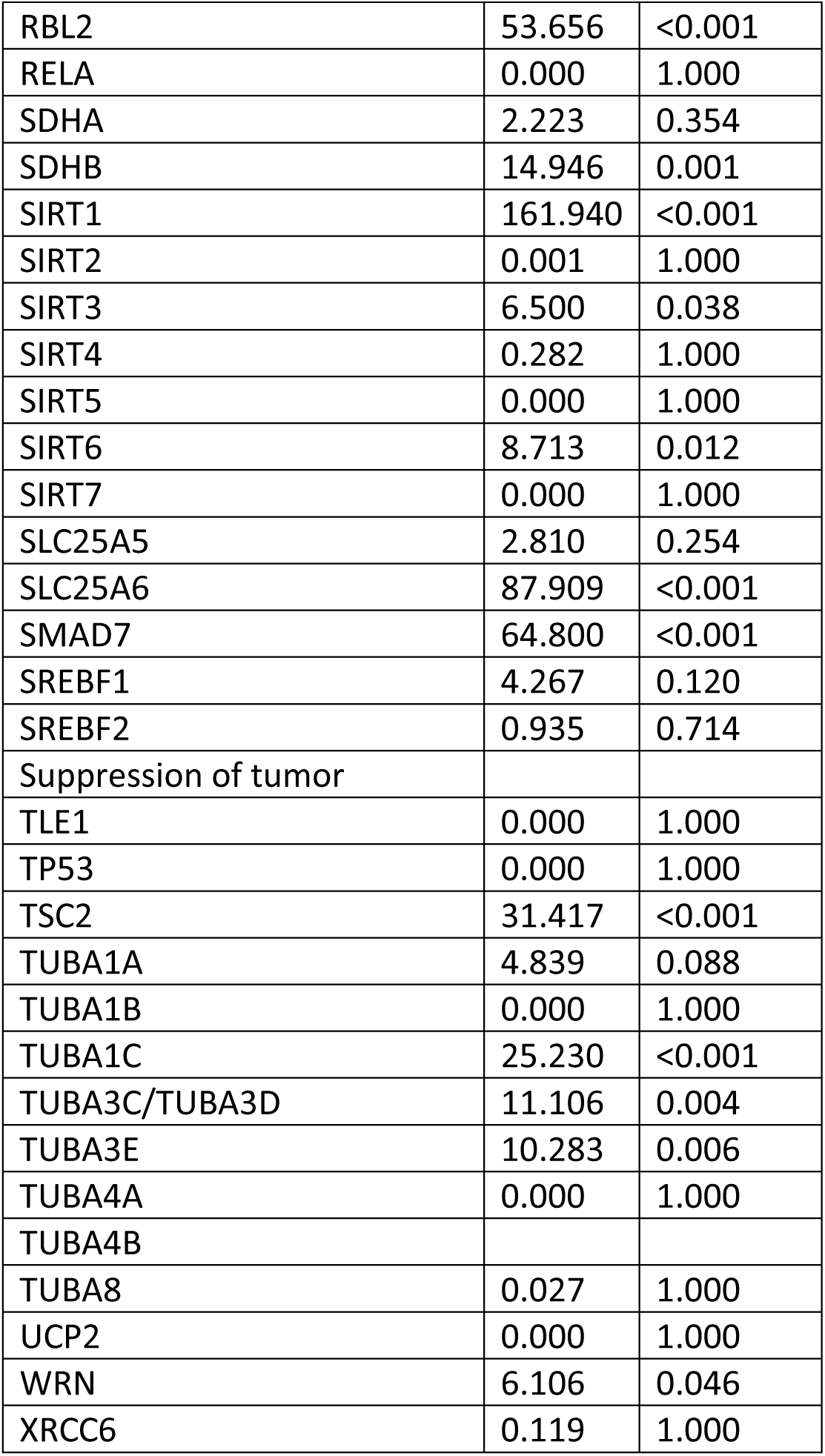
SIRT signaling pathway overlaid with the likelihood-ratio test statistics (2ΔL) between the positive and neutral branch-models as a measure to visualize a pathway effect.

**Table S6.** P53 signaling pathway overlaid with the likelihood-ratio test statistics (2ΔL) between the positive and neutral branch-models as a measure to visualize a pathway effect.

| Symbol | $2\Delta L$ | *p*-value |
| --- | --- | --- |
| 14-3-3σ | 0.000 | 1.000 |
| AKT |  |  |
| Angiogenesis |  |  |
| Apaf1 | 1.169 | 0.625 |
| Apoptosis |  |  |
| ASPP |  |  |
| ATM | 2.078 | 0.382 |
| ATR | 0.065 | 1.000 |
| Autophagy |  |  |
| BAI1 | 2.103 | 0.378 |
| BAX | 0.000 | 1.000 |
| Bcl-2 | 0.077 | 1.000 |
| Bcl-xL | 0.000 | 1.000 |
| Brca1 |  |  |
| CABC1 |  |  |
| Caspase 6 |  |  |
| CDK2 | 0.347 | 1.000 |
| CDK2-Cyclin D1 |  |  |
| CDK4 | 0.000 | 1.000 |
| CDK4-Cyclin D2 |  |  |
| Cell cycle arrest |  |  |
| Cell cycle progression |  |  |
| Cell survival |  |  |
| Chk1 | 0.000 | 1.000 |
| Chk2 | 0.001 | 1.000 |
| c-Jun | 2.366 | 0.327 |
| CK1δ |  |  |
| Cyclin D1 | 0.014 | 1.000 |
| Cyclin D2 | 0.010 | 1.000 |
| CyclinG | 13.155 | 0.001 |
| CyclinK |  |  |
| DNA damage |  |  |
| DNA repair |  |  |
| DNA-PK |  |  |
| DR4/5 |  |  |
| DRAM |  |  |
| E2F1 | 1.341 | 0.566 |
| E2f1-Rb |  |  |
| FAS | 1.763 | 0.450 |
| GADD45 |  |  |
| Glycolysis |  |  |
| GML |  |  |
| Gsk3β | 55.087 | <0.001 |
| HDAC |  |  |
| HDAC9 |  |  |
| HIF1A | 0.000 | 1.000 |
| HIPK2 |  |  |
| Hypoxia |  |  |
| JMY |  |  |
| Jmy-p300 |  |  |
| JNK1 | 0.001 | 1.000 |
| Maspin | 0.223 | 1.000 |
| MDM2 | 0.000 | 1.000 |
| MDM4 | 0.001 | 1.000 |
| Mitochondrial respiration |  |  |
| NOXA | 9.495 | 0.008 |
| Nucleostemin |  |  |
| p19arf | 23.352 | <0.001 |
| p21Cip1 | 0.000 | 1.000 |
| p300 |  |  |
| p38 MAPK |  |  |
| p48 |  |  |
| p53 | 0.000 | 1.000 |
| p53AIP1 |  |  |
| p53R2 | 0.018 | 1.000 |
| p63 |  |  |
| p73 | 0.564 | 0.915 |
| PAI-1 |  |  |
| PCAF |  |  |
| PCNA |  |  |
| PERP | 22.105 | <0.001 |
| PI3K |  |  |
| PIAS1 |  |  |
| PIDD | 0.001 | 1.000 |
| PIG3 | 18.848 | <0.001 |
| PML |  |  |
| PTEN | 0.019 | 1.000 |
| PUMA | 33.998 | <0.001 |
| Rb | 61.629 | <0.001 |
| Reprimo | 0.000 | 1.000 |
| SCO2 |  |  |
| Senescence |  |  |
| SIRT | 161.940 | <0.001 |
| Slug |  |  |
| STAG1 |  |  |
| Survivin | 7.876 | 0.019 |
| Teap |  |  |
| TIGAR |  |  |
| TOPBP1 |  |  |
| TRAP220 |  |  |
| TRIM29 |  |  |
| TSP1 | 0.016 | 1.000 |
| Tumor suppression |  |  |
| UCN-01 |  |  |
| WT1 |  |  |
| ZAC1 |  |  |
| β-catenin | 0.001 | 1.000 |

**Table S7.** Insulin signaling pathway overlaid with the likelihood-ratio test statistics (2ΔL) between the positive and neutral branch-models as a measure to visualize a pathway effect.

| Symbol | $2\Delta L$ | $p$-value |
| --- | --- | --- |
| AKT |  |  |
| BCL-XL | 0.000 | 1.000 |
| c-FOS | 0.000 | 1.000 |
| c-JUN | 2.366 | 0.327 |
| c-Raf | 105.387 | <0.001 |
| CCK2R | 0.000 | 1.000 |
| CEBPβ | 0.000 | 1.000 |
| Cell proliferation |  |  |
| CIS | 0.000 | 1.000 |
| ERK1/2 |  |  |
| Gαq | 0.000 | 1.000 |
| GAST | 2.472 | 0.311 |
| GRB2 | 50.205 | <0.001 |
| IL-6 | 48.957 | <0.001 |
| JAK |  |  |
| JAK2 | 0.043 | 1.000 |
| LEP | 0.000 | 1.000 |
| LEPR | 0.000 | 1.000 |
| MEK1/2 |  |  |
| mTOR | 41.010 | <0.001 |
| NFκB | | |
| p21Cip1 | 0.000 | 1.000 |
| PI3K |  |  |
| PIAS |  |  |
| PTP1B | 14.851 | 0.001 |
| Ras |  |  |
| SHC | 0.000 | 1.000 |
| SHP1 | 0.000 | 1.000 |
| SHP2 | 9.205 | 0.009 |
| SOCS |  |  |
| SOS |  |  |
| STAT |  |  |
| Stat dimer |  |  |
| STAT3 | 2.391 | 0.323 |
| Stat3 dimer |  |  |

